# Motor and cognitive deficits limit the ability to flexibly modulate spatiotemporal gait features in older adults with mild cognitive impairment

**DOI:** 10.1101/2022.09.09.507368

**Authors:** Michael C. Rosenberg, Alexandra Slusarenko, Ke Cao, J. Lucas McKay, Laura Emmery, Trisha M. Kesar, Madeleine E. Hackney

**Affiliations:** Neuromechanics Laboratory, Department of Biomedical Engineering, Emory University & Georgia Institute of Technology, Atlanta, GA, USA; College of Arts and Sciences, Emory University, Atlanta, GA, USA; Department of Medicine, Division of Geriatrics and Gerontology, Emory University School of Medicine, Atlanta, GA, USA; Department of Neurology, Emory University School of Medicine, Atlanta, GA, USA; Department of Biomedical Informatics, Emory University School of Medicine, Atlanta, GA, USA; Department of Music, Emory University College of Arts and Sciences, Atlanta, GA, USA; Department of Rehabilitation Medicine, Emory University School of Medicine, Atlanta, GA, USA; Emory University School of Nursing, Atlanta, GA, USA; Atlanta VA Center for Visual & Neurocognitive Rehabilitation, Atlanta, GA, USA; Birmingham/Atlanta VA Geriatric Research Education and Clinical Center, Atlanta, GA, USA

**Keywords:** mild cognitive impairment, dance therapy, gait analysis, biomechanics, music, rhythm, gait modifications

## Abstract

**Introduction:** Dance-based therapies are an emerging form of movement therapy aiming to improve motor and cognitive function in older adults with mild cognitive impairments (MCIs). Despite promising effects of dance-based therapies on function, it remains unclear how age-related declines in motor and cognitive function affect movement capacity and influence which movements and rhythms maximize dance therapy efficacy. Here, we evaluated the effects of age and MCI on the ability to accurately modulate spatial (*i*.*e*., joint kinematics), temporal (*i*.*e*., step timing), and spatiotemporal features of gait to achieve spatial and temporal targets during walking.

**Methods:** We developed novel rhythmic movement sequences - nine spatial, nine temporal, and four spatiotemporal - that deviated from typical spatial and temporal features of walking. Healthy young adults (HYA), healthy older adults (HOA), and adults with MCI were trained on each gait modification before performing the modification overground, with kinematic data recorded using wearable sensors.

**Results:** HOA performed spatial (p = 0.010) and spatiotemporal (p = 0.048) gait modifications less accurately than HYA. Individuals with MCI performed spatiotemporal gait modifications less accurately than HOA (p = 0.017). Spatial modifications to the swing phase of gait (p = 0.006, Cohen’s *d* = -1.3), and four- and six-step *Duple* rhythms during temporal modifications (p ≤ 0.030, Cohen’s *d* ≥ 0.9) elicited the largest differences in gait performance in HYA vs. HOA and HOA vs. MCI, respectively.

**Discussion:** These findings suggest that age-related declines in strength and balance reduce the ability to accurately modulate spatial gait features, while declines in working memory in individuals with MCI may reduce the ability to perform longer temporal gait modification sequences.

Differences in rhythmic movement sequence performance highlight motor and cognitive factors potentially underlying deficits in gait modulation capacity, which may guide therapy personalization and provide more sensitive indices to track intervention efficacy.

## Introduction

Dance-based therapy, which purposefully sets movements to music, is an emerging form of movement therapy for individuals with mild cognitive impairments (MCIs), a precursor to dementia or Alzheimer’s involving reduced attention, executive function, spatial cognition, and working memory, that elicits improvements in cognitive function in individuals with MCI (Gauthier et al., 2006; Lazarou et al., 2017; Sanford, 2017; Zhu et al., 2018; Zhu et al., 2020). Dance-based therapy is a logical approach for improving cognitive function, as exercise or general physical activity are among the few interventions shown to improve cognitive function in individuals with MCI (Geda et al., 2010; Thom and Clare, 2011). Specifically, Thom and Clare (2011) noted that physical exercise programs that target functional tasks, aerobic endurance, and balance are likely needed to maximize the functional benefits of therapy, all of which are challenged during dance (Thom and Clare, 2011). Currently, it is unclear how to personalize dance-based therapy parameters for individuals with MCI. Studies of dance-based therapy use different music, dance types, and study designs (Hackney and Earhart, 2009; McKee and Hackney, 2013; Lazarou et al., 2017; Zhu et al., 2020). Improving our understanding of the relationships between individual-specific deficits associated with MCI and the ability to perform different movements during dance-based therapy is needed to inform the objective design of personalized dance-based therapies for individuals with MCI.

While a major goal of dance-based therapies is to maximize the motor and cognitive benefits of moving to music, *how* to do so for a given population needs better clarity. Music type and tempo, and the dance type or steps are basic parameters that can influence the difficulty of performing the therapy routine. Therapies often rely on trial-and-error to refine a protocol, rather than being grounded in a principled data-driven understanding of myriad factors that may impact dance-therapy performance, including individual differences in motor control and cognitive function (McKee and Hackney, 2013; Lazarou et al., 2017; Zhu et al., 2018). Selecting an appropriate level of challenge is fundamental to maximizing the motor and cognitive effects of therapy (Guadagnoli and Lee, 2004). Given the heterogeneity of cognitive deficits in MCI (Gauthier et al., 2006) and age-related declines in motor function (Booth et al., 1994), understanding how both motor and cognitive deficits impact the ability to perform challenging rhythmic movement patterns may both enable objective personalization of dance-based therapy parameters and provide mechanistic insights into the impacts of MCI on mobility.

Though not yet quantified in the context of dance-based therapy, dance can be considered a series of Rhythmic Movement Sequences (RMS): rhythmic patterns involving modifications to spatial and temporal components of normal stepping (Rallis et al., 2018). Importantly, spatial and temporal features of movement correspond to, and may challenge, distinct neural representations (*e*.*g*., the premotor cortex) and multiple aspects of biomechanics function (*e*.*g*., strength, balance, joint coordination) (Wilson and Kwon, 2008; Kornysheva and Diedrichsen, 2014). A central premise of this work is that evaluating the capacity for modulating spatial and temporal movement features will shed light on motor and cognitive mechanisms that may inform future therapy individualization and enhance the efficacy of dance and music-based movement therapy in individuals with MCI. Spatial modifications of movement during dance involve altering the magnitude and coordination of hip, knee, and ankle kinematics (Ivanenko et al., 2005; Wilson and Kwon, 2008). Voluntarily altering coordination relative to highly stereotyped patterns during locomotion requires the motor capacity to achieve joint excursions and increased cortical demands (Ivanenko et al., 2005; Cohen et al., 2016; Hortobágyi et al., 2016; Rucco et al., 2017). Consequently, both motor and cognitive deficits may impact individuals’ abilities to perform modifications to the spatial components of movement.

Muscle function and motor control change with age, and correspond to reductions in overall motor function in older adults (Booth et al., 1994; Schloemer et al., 2017). Therefore, age-related deficits in motor function may impact individuals’ abilities to modify spatial components of movement.

Because spatial modifications also require mapping perceived biomechanical targets to motor actions, we expect these modifications to challenge aspects of cognitive function that are impaired in individuals with MCI, such as spatial attention, executive function, and spatial cognition (Gauthier et al., 2006; Rucco et al., 2017). However, how these motor and cognitive deficits impact individuals’ abilities to voluntarily modulate spatial aspects of movement remains unclear.

Temporal modifications of movement involve altering step timing to produce rhythmic sequences of steps, which may challenge cognitive function (Kornysheva and Diedrichsen, 2014). During dance therapy, temporal modifications can be elicited by altering the music’s rhythm and meter, two fundamental components of music (Hackney et al., 2007a; Hackney and Earhart, 2009). In the context of stepping or walking, rhythm defines the pattern of temporal stimuli, such as taking two quick (q) steps followed by two slow (S) steps. Meter is a perceived quantity that individuals anticipate and to which they entrain their movements. Unimpaired walking typically involves evenly recurring steps that are approximately rhythmically and metrically constant, such that modifying temporal features during RMS would represent a deviation from normal gait rhythm and meter.

Because temporal modifications require attention and the ability to perceive rhythmic cues and map them to motor actions, we expect it to challenge aspects of cognitive function that are impaired in individuals with MCI, such as attention and working memory (Gauthier et al., 2006; Grahn, 2012; Sanford, 2017). Because temporal modifications can be performed with normal joint coordination patterns, they are less likely to challenge motor function than spatial modifications. Therefore, we expect spatial modifications to movement during RMS to challenge both motor and cognitive function, while temporal modifications should primarily challenge cognitive function. However, the extent to which age-related motor and cognitive deficits impact the ability to modulate spatial and temporal aspects of movement is unclear.

In this study, we leveraged biomechanical analysis, prototypical dance movements, and music theory to systematically investigate the effects of age-related differences in motor and cognitive function on the ability to perform spatial and temporal modifications to walking in individuals with MCI. We hypothesized that differences in motor and cognitive function would impact individuals’ abilities to accurately modulate spatial and temporal features of walking, similar to those performed during dance-based therapies. Specifically, we predicted that, when performing RMS gait modifications of varying difficulty, healthy older adults (HOA) would perform spatial and spatiotemporal gait modifications significantly less accurately than healthy young adults (HYA), suggesting an effect associated with aging-related differences in motor function. Similarly, we predicted that individuals with MCI would perform temporal and spatiotemporal RMS less accurately than HOA, suggesting an effect associated with impaired cognitive function.

## Materials and Methods

This study was approved by the Emory University Institutional Review Board (STUDY00003507). All participants gave written informed consent prior to participation.

### Study design and visits

We performed an observational cross-sectional study in healthy younger and older adults, and older adults with MCI. All outcomes were collected during a single 2-4 hour study visit for each participant. Participants who were unable to complete the protocol in a single visit completed it during one additional visit within one week.

### Participants

A total of 37 participants were recruited for the study, including younger adults (HYA, N=13), older adults (HOA, N=12), and older adults with MCI (N=12; **Error! Reference source not found**.). One HYA participant failed to complete the protocol and was excluded from analysis. For all participants, inclusion criteria included the ability to walk 20m without an assistive device, completion of at least 6 years of education or good work history, no hospitalizations in the last 60 days, and proficiency in the English language. HYA participants ages 18-35 years were included in this study and HOA participants ages 55 years and older were included. Additional inclusion criteria for participants with MCI included amnestic MCI defined using the Alzheimer’s Disease Neuroimaging Initiative criteria and reduced executive function, working memory, and spatial cognition according to standard clinical assessments (Phillips et al., 2001; Vandierendonck et al., 2004; Mueller et al., 2005; Bowie and Harvey, 2006; Hackney et al., 2013).

### Motor and cognitive assessments

To characterize participants’ motor and cognitive function, trained personnel administered the following motor and cognitive assessments: Montreal Cognitive Assessment (MoCA), Reverse Corsi Blocks test, Tower of London test, Trail Making Test, Four-Square Step Test, Body Position Spatial Task, simple, manual, and cognitive Timed Up-and-Go (TUG), and the 30-second Chair Stand (Lundin-Olsson et al., 1998; Jones et al., 1999; Morris et al., 2001; Phillips et al., 2001; Dite and Temple, 2002; Vandierendonck et al., 2004; Nasreddine et al., 2005; Bowie and Harvey, 2006; Hackney et al., 2013). The average (± 1SD) scores for each clinical assessment and corresponding p-values denoting the probability of statistical differences in each score were computed using the software package R (**Error! Reference source not found**.). Older adults and individuals with MCI were well-matched in age and sex. As expected, the MCI participants trended toward worse performance on the tests of motor and cognitive function. HYA participants performed better than HOA or MCI participants on all assessments of motor and cognitive function. For comparison to the tempi of temporal and spatiotemporal gait modifications, we also recorded participants’ stride frequencies at their preferred walking speeds.

### RMS gait modifications

Participants performed normal walking at their self-selected and fast speeds, and 22 RMS consisting of 9 spatial, 9 temporal, and 4 spatiotemporal gait modifications, described below. Participants were randomly assigned to perform either the spatial or temporal modifications first. Spatiotemporal modifications always occurred after spatial and temporal modification trials. Experimental conditions within each sub-group (spatial, temporal, or spatiotemporal) were randomized.

We defined 64 spatial, temporal, and spatiotemporal gait modifications, constituting a movement library, from which we selected 9 spatial, 9 temporal, and 4 spatiotemporal gait modifications of varying levels of difficulty. The selected biomechanical (*i*.*e*., spatial) and temporal features of each modification are described in **Error! Reference source not found.A**. The figure shows the kinematic configuration of each movement and lists the target variables. Briefly, *spatial* modifications altered the magnitude of hip, knee, and ankle movements and their coordination relative to normal walking (Ivanenko et al., 2005). We selected three stance-phase modifications (ST), three swing-phase modifications (SW), and three combined modifications in both the swing and stance phases (SWST). Participants were provided target joint angles for each modification, such as flexing the hip and knee to 90 degrees during swing; and watched a video of an expert dancer performing the movements. During ST and SW trials, participants modified stance and swing-phase gait kinematics, respectively, in both legs asymmetrically. During SWST trials, participants modified both swing and stance only in the left leg.

**Figure 1:**
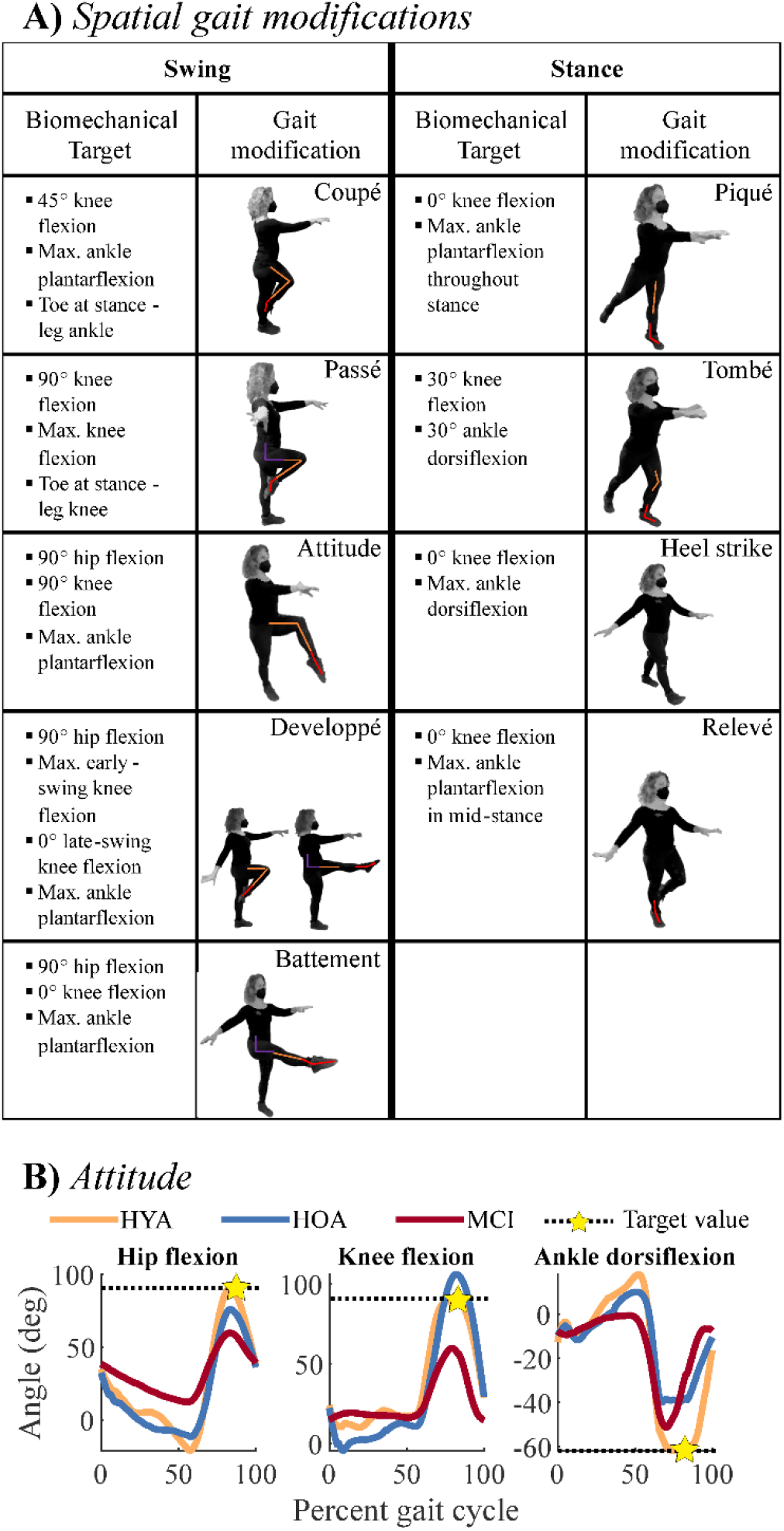
Descriptions of subcomponents of the spatial gait modifications. or rhythmic movement sequences (RMS) evaluated in the current study. A) Spatial gait modifications were derived from ballet steps, noted in the boxes containing movement depictions. The left two columns correspond to swing-phase modifications, while the right two columns correspond to stance-phase modifications. Each depiction shows the modification where biomechanical targets were applied. Colored lines on each person denote biomechanical targets at the hip (purple), knee (orange), and ankle (red) kinematics that were used to evaluate movement performance. Bulleted lists describe the biomechanical targets for each gait modification, which were used to quantify movement performance. Bullets denote the biomechanical targets for each gait modification. Modifications were performed in pairs during spatial trials. B) Example of biomechanical targets for the *Attitude* gait modification. Black dashed lines denote biomechanical target values and gold stars denote where the target value was estimated. For example, during the *Attitude* modification, the swing-leg hip (left) and knee (middle) should be flexed to 90 degrees, and the ankle (right) should be maximally plantarflexed. Colored lines denote exemplary HYA (orange), HOA (blue), and MCI (red) participants. These participants highlight reduced RMS performance at the hip and knee in MCI and at the ankle in HOA and MCI

*Temporal* modifications altered the rhythmic patterns of movement, consisting of two, four, or six steps taken at half-beat (“and;” denoted *&*), one-beat (“quick;” denoted *q*), and two-beat (“slow;” denoted *S*) intervals (**Error! Reference source not found**.). The music notation used in **Error! Reference source not found**. indicates the sequence and duration of steps, with closed (black) notes corresponding to a step spanning one beat and open (white) notes denoting steps spanning two beats. Performance was evaluated by each participant’s ability to take steps matching the beat and in the correct sequence of quick and slow steps.

**Figure 2:**
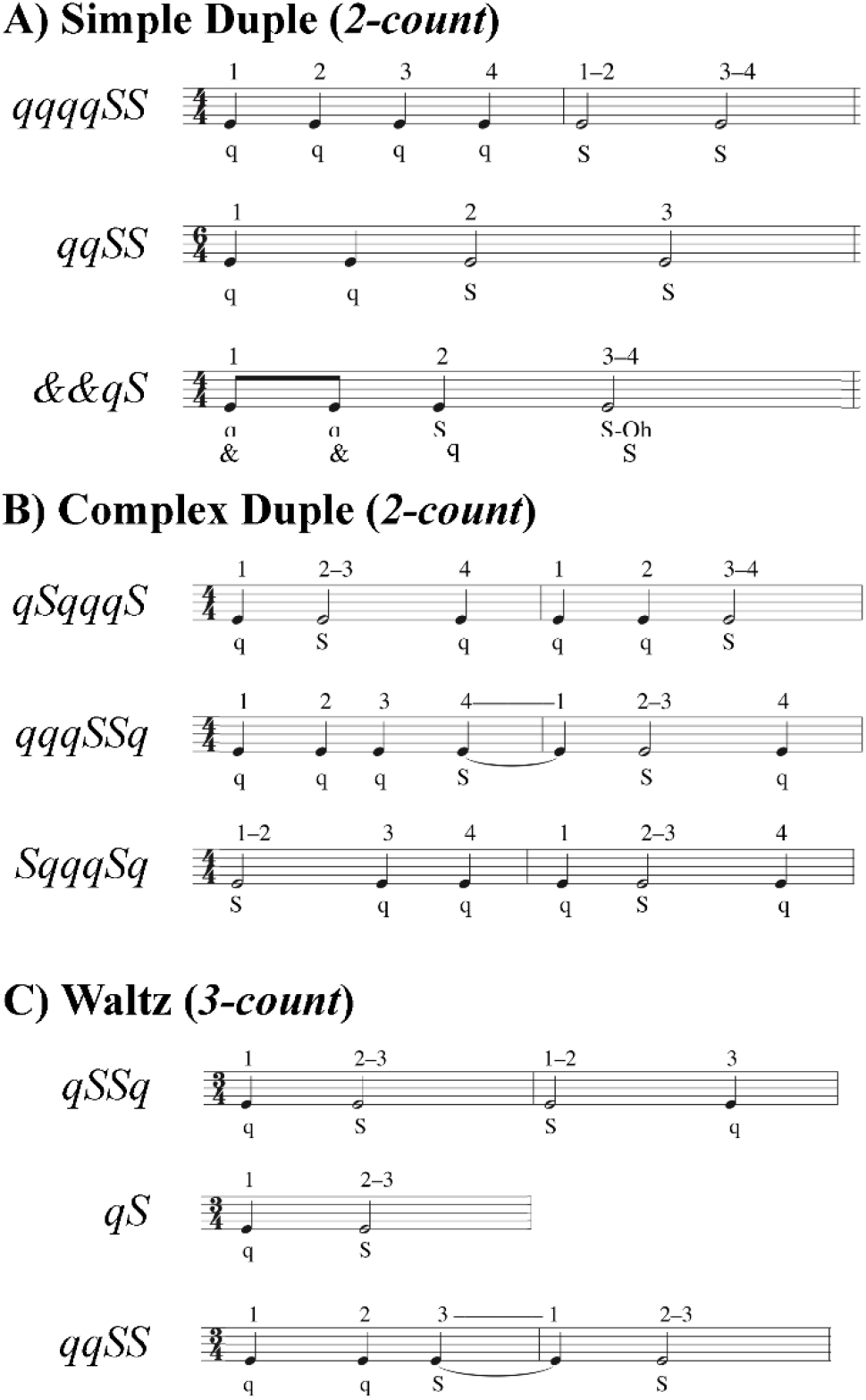
Step sequences for temporal and spatiotemporal gait modifications. Step sequences consisted of 2, 4, or 6 steps. Each sequence specified a combination of very quick (*&* = half-beat), quick (*q* = one beat), and slow (*S* = two beats) steps. The numbers above the musical bars show the beat count for each rhythm. A) Simple duple sequences used a two-count rhythm in which the strong beat was the downbeat. B) Complex duple sequences also used a two-count rhythm, but the strong beat was not the downbeat. C) Waltz sequences used a three-count rhythm, such that sequences spanned either one or two measures.

Temporal modifications were split into three sub-groups: three simple duple, three complex duple, and three waltz modifications. *Duple* music (**Error! Reference source not found**.A & B) had a tempo of 100 beats per minute (bpm) for all participants, while Waltz music (**Error! Reference source not found**.C) had a tempo of 134 bpm for HYA but was reduced to 104 bpm for the HOA and MCI groups. The faster Waltz tempo for the HYA group was due to an error in coding the tempi for the first 8 participants. We corrected the tempo for the HOA and MCI to ensure that all participants could perform the temporal modifications. We used the same tempi across all participants, rather than personalizing the tempi to each participant’s preferred cadence to avoid giving participants different amounts of time to process and remember auditory input, which could alter cognitive demands for each participant (Gauthier et al., 2006; Persad et al., 2008; Montero-Odasso et al., 2012). The *simple duple* meter has a two-count grouping (1-2) where the downbeat is the strong beat. In more *complex duple* meters, the emphasized beat falls on the weak beat, which defies Western music listeners’ expectations. Consequently, we expected the complex duple to be more challenging than the simple duple. The *waltz* is in a compound duple meter (a fast two-count where each is subdivided into three sub-beats; 1-2-3, 1-2-3) or simple triple (three-count; 1-2-3) meter. *Waltz* meters, unlike *simple* and *complex duple* meters, in which synchronization of movement to music is easier because the same foot always lands on the downbeat, the three-count of *waltz* rhythms causes the foot landing on the downbeat to alternate every measure (three steps).

Consequently, individuals accustomed to Western music traditions may have greater difficulty synchronizing movements to the triple meter of *waltz* than to duple meters. The movements within each sub-group varied in difficulty. During duple trials, participants were cued by a simplified *Libertango* (Astor Piazzolla, 1974), with superfluous musical accents and cues eliminated to improve the ability of participants with limited musical experience to identify temporal patterns in the music. During waltz trials, participants were cued by a modified version of *Shostakovich Waltz No. 2* (Dmitri Shostakovich, 1938). These musical pieces are much more complex than a metronome beat used in locomotion studies, as the attention required to dissociate meter and rhythm from superfluous auditory information can alter individuals’ abilities to entrain to the tempo (Styns et al., 2007; Moumdjian et al., 2019). For each modification, participants were provided a sequence of “quick” and “slow” beats and practiced the sequence by following an instructional video: first clapping, then tapping the foot, then shifting weight between the legs, then stepping in place, then walking (Hackney and Earhart, 2010).

*Spatiotemporal* modifications combined spatial and temporal modifications of varying difficulty (**Error! Reference source not found**. & **Error! Reference source not found**.). We selected these movements to vary the relative motor and cognitive challenges of the movements. Participants were allowed to review spatial and temporal instructional videos if needed. Following instructions for all modifications, the assessors would briefly practice (3-5 sequences) the modifications with the participant to confirm understanding. Data were collected immediately after understanding was confirmed. Names and descriptions of all gait modifications can be found in the *MovementLibrary*.*csv* and *DataDictionary*.*csv*, with reference to **Error! Reference source not found**. & **Error! Reference source not found**..

During each trial, participants wore 15 inertial measurement units (IMUs) in a standard configuration (APDM, Inc., Portland, USA). We used validated commercial software associated with the IMUs to estimate lower-limb hip, knee, and ankle flexion angles over the gait cycle (Mancini and Horak, 2016; Washabaugh et al., 2017). Participants began each trial in quiet stance until hearing a tone, then began walking in a modified gait pattern defined by the trial’s spatial, temporal, or spatiotemporal RMS. When physically capable, participants walked overground along an 11m walkway four times for each trial, providing 15-30 (spatial) or 15-60 (temporal) strides for each trial. For participants with reduced physical capacity, we iteratively reduced the number of walkway lengths traveled from four lengths down to one length based on each participant’s physical capacity.

We ensured that each participant performed at least 15 strides for spatial modifications and 10 sequences for temporal and spatiotemporal modifications. However, all participants performed the same gait modifications, regardless of the distance walked. Participants were allowed to rest as needed and were provided snacks and water between trials. All assessments were administered by trained personnel.

### RMS performance

For each modification, we defined biomechanical (spatial and spatiotemporal trials) and temporal (temporal and spatiotemporal trials) targets, defined *a priori* in our movement library (**Error! Reference source not found.A**). Biomechanical targets matched those provided during instruction. Temporal targets corresponded to the step sequence of quick and slow steps. RMS performance was defined as a percent error relative to biomechanical and temporal targets: the error relative to each target value, normalized by the magnitude of the corresponding target (**Error! Reference source not found.B**). For example, the hip flexion angle target for the *attitude* movement, the star in **Error! Reference source not found.B**, is 90 degrees during the swing phase of gait. In swing, the knee should be flexed to 90 degrees, and the ankle maximally plantarflexed. The targets for each spatial modification are listed in **Error! Reference source not found.A**. Therefore, RMS performance reflected the accuracy with which each participant performed a gait modification, with lower percent error reflecting more accurate (*i*.*e*., better) performance. For each trial and participant, we averaged across each trial’s biomechanical and temporal targets (Equation <u>(1</u>). For each gait modification class (*spatial, temporal, spatiotemporal*), we computed the error the *i*^*th*^ participant made on the *j*^*th*^ trial of the *k*^*th*^ target variable (e.g., peak knee flexion angle) as follows:

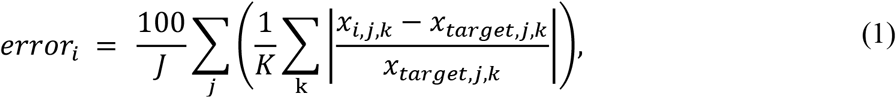

where *x*_*i,j,k*_ denotes the *i*^*th*^ participant’s outcome for the *j*^*th*^ trial and *k*^*th*^ target variable (*e*.*g*., peak knee flexion angle), and is a vector of outcomes over all available strides for that trial. The variable *x*_*target,j,k*_ denotes the value of the *j*^*th*^ trial and *k*^*th*^ target variable. The variable *J* represents the number of trials included in each modification class and *K* represents the number of pre-specified target variables for each gait modification (*spatial*: 9 modifications, 2-3 target variables per trial; *temporal*: 9 modifications, 1 target variable per trial; *spatiotemporal*: 4 modifications, 3-4 target variables per trial). Percent error for each target (multiple targets per trial; see **Error! Reference source not found**. & **Error! Reference source not found**.) was averaged to compute a single error value for the trial. When multiple trials were included in a single analysis (*e*.*g*., all *spatial* trials), the percent errors were averaged across trials. We computed RMS performance outcomes using all available data from each trial, regardless of trial length.

### Statistical analysis

Group differences in demographic and clinical characteristics were assessed with omnibus analysis of variance (ANOVA) and Chi-squared test, as appropriate. Significant results (α = 0.05) were followed by post-hoc independent sample t-tests for differences between the HYA and HOA groups, and between the HOA and MCI groups. Based on our hypotheses, we only compared the HYA and HOA groups, and the HOA and MCI groups.

We examined differences in RMS performance across groups using separate fixed effects linear models for each outcome variable. To test for group differences in RMS performance between the HYA and HOA groups, and between HOA and MCI groups, we used linear models containing indicator variables denoting the HYA and MCI groups, with HOA as the reference group (Equation (2). Significant coefficients (Wald Tests, α = 0.05) corresponding to the HYA (β _*HYA*_) or MCI variables (β _*MCI*_) would indicate significant differences in performance between the HOA group and the HYA or MCI groups, respectively. Separate linear models were fit for each of the three gait modification classes (*spatial, temporal, spatiotemporal*). The sample size for each model was N = 36. Because we were not interested in comparisons between HYA and MCI, we parameterized linear models so that the coefficients would directly test the contrasts of interest.

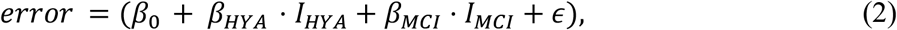

Where *error* is a 36×1 vector with one sample for each participant. The variable β _0_ denotes the average percent error of the HOA group, β _*HYA*_ and β _*MCI*_ are coefficients corresponding to the magnitude of differences between the HYA and HOA groups and the HOA and MCI groups, respectively. Variables *I*_*HYA*_ and *I*_*MCI*_ are indicator variables corresponding to the respective coefficients. The variable Є is an error term. Linear models were fit using *fitlm* in MATLAB 2021b (Mathworks Ltd, Natick, USA), which estimates coefficients using least-squares regression and determines the coefficients’ p-values using Wald tests. To determine whether specific types of gait modifications were differentiated between groups, we repeated the above analysis for gait modification subclasses (*e*.*g*., *swing, simple duple*) and individual gait modifications (*e*.*g*., *attitude-developpe, qqqqSS*; **Error! Reference source not found**.A & **Error! Reference source not found**.A).

To determine which movements best distinguished between groups, we computed effects sizes (Cohen’s *d*; vs. HOA) and the average difference (Δ vs. HOA) for each gait modification class (*spatial, temporal, spatiotemporal*). To determine whether specific gait modifications were differentiated between groups, we repeated the above analysis for gait modification subclasses (*e*.*g*., *swing, simple duple*).

As an exploratory investigation to identify potential correlates of RMS performance, we computed separate Pearson correlations between each of the clinical assessments of motor and cognitive function (**Error! Reference source not found**.) against *spatial, temporal*, and *spatiotemporal* gait modification performance, with N=36 samples per correlation.

## Results

### Study compliance and participants

Data were collected in 37 participants (12 per group, plus one additional HYA participant). The additional HYA participant was unable to follow directions when performing rhythmic movement sequences and was excluded from analysis. One MCI participant failed to complete the protocol due to fatigue and completed the protocol in a follow-up visit. One MCI participant was only able to complete temporal trials due to fatigue and balance concerns. Across the 36 participants included in analysis, only 19 trials (less than 2% of all trials) were omitted due to sensor errors not detected during the experiment. Most participants completed the full four walkway lengths of the RMS protocol. Across groups, 24 participants completed four walkway lengths, 10 participants completed two-to-three lengths (5 HOA, 5 MCI), and only two participants (1 HOA, 1 MCI) completed a single walkway length.

Demographic characteristics for each participant group are shown in **Error! Reference source not found**.. Participants exhibited similar demographic characteristics, as expected. Clinical characteristics for each participant group are shown in **Error! Reference source not found**..

Individuals with an MCI diagnosis trended toward worse performance than HOA on the Trail Making Test (p = 0.099; independent-samples t-tests), cognitive (p = 0.084), and manual (p = 0.067) TUG tests. HOA performed worse than HYA on the Reverse Corsi Blocks (p = 0.003), Body Position Spatial Task (p = 0.028), and the Four-Square Step Test (p = 0.002). All three groups walked with similar cadences.

### RMS performance was reduced with age

When averaging across gait modifications within each modification class (*spatial, temporal, spatiotemporal*), regression analysis revealed that the HYA group performed *spatial* gait modifications significantly more accurately than the HOA group (p = 0.010; *d* = 1.1, Δ = 4.0%) (**Error! Reference source not found**.). HYA participants also performed *spatiotemporal* gait modifications more accurately than HOA participants (p = 0.048; *d* = 0.8, Δ = 2.1%). Conversely, HYA participants did not perform *temporal* gait modifications significantly more accurately than HOA participants.

**Figure 3:**
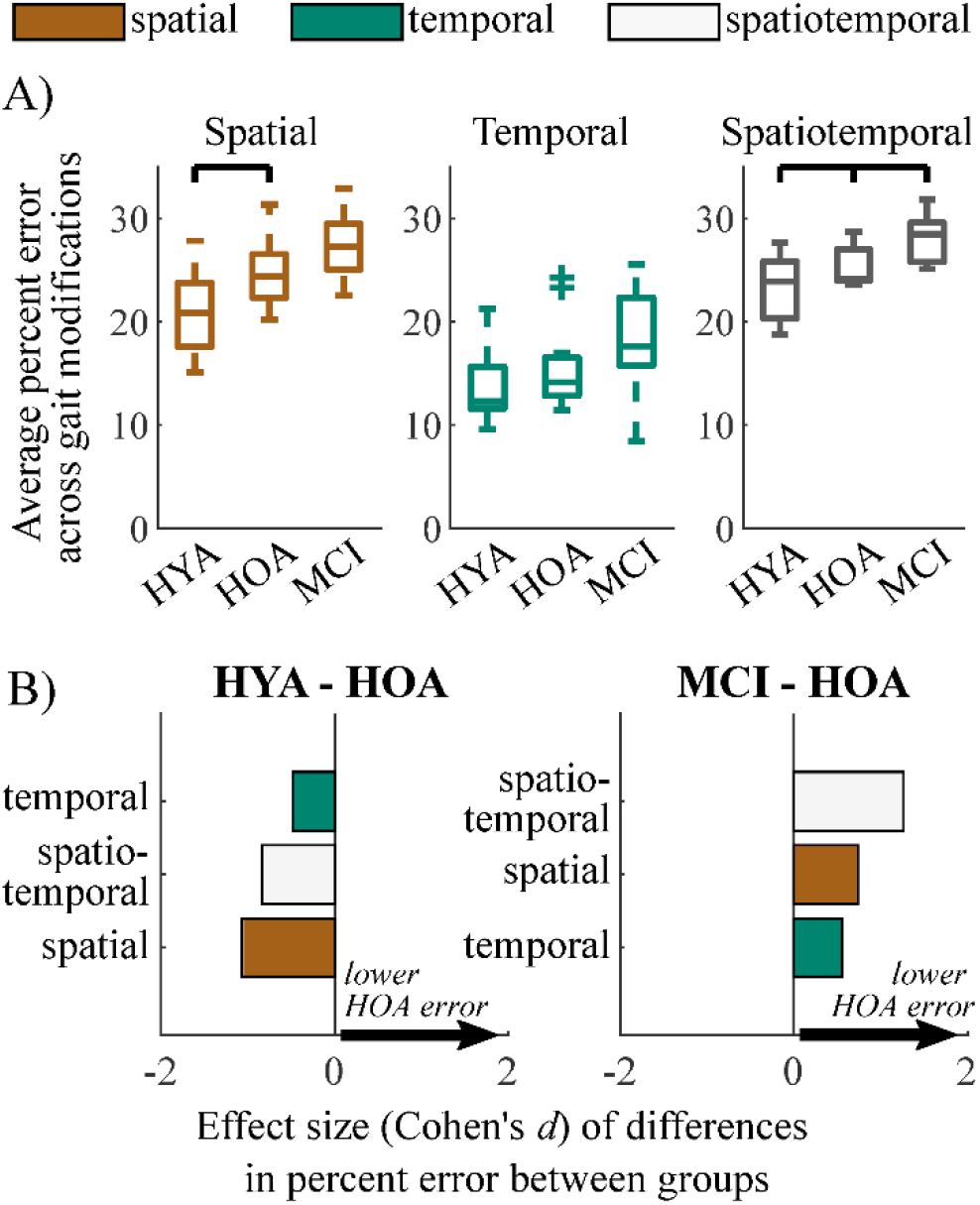
Summary of group performance on gait modifications. A) Boxplots showing group performance on spatial, temporal, and spatiotemporal trials. Each sample consists of the average percent error relative to target values across nine gait modifications for spatial and temporal trials and four gait modifications for spatiotemporal trials. B) Effect sizes of differences in performance (Cohen’s *d*) between HYA and HOA groups (left) and HOA and MCI groups (right) on spatial (orange), temporal (green), and spatiotemporal (white) gait modifications. Positive values (arrows) imply that the HOA group exhibited lower error on gait modifications. We did not test for differences between the HYA and MCI groups.

Within movement subclasses (*e*.*g*., *swing, simple-duple*), differences in spatial performance between the HYA and HOA groups were largest in modifications to the *swing* phase of gait (p = 0.006, *d* = -1.3, Δ = 5.8%) (**Error! Reference source not found**.A & B). Differences in gait modification performance in both the *stance* phase (p = 0.074, *d* < 0.8, Δ = 2.2%) and the combined *swing-stance* modifications (p = 0.054, *d* < 0.9, Δ = 2.8%) had smaller effects that did not reach significance. In particular, deficits in gait modification accuracy in HOA were largest in *swing*-phase modifications that involved large hip flexion ranges of motion (*i*.*e*., *attitude, developpé*, and *battement*; see modification descriptions in **Error! Reference source not found.A** and supplemental file *MovementLibrary*.*csv*), with large effect sizes (p < 0.001; *d* > 0.9; Δ = 4.5%). These differences are exemplified by the *attitude-developpé* modification shown in Error! Reference source not found.**C**.

**Figure 4:**
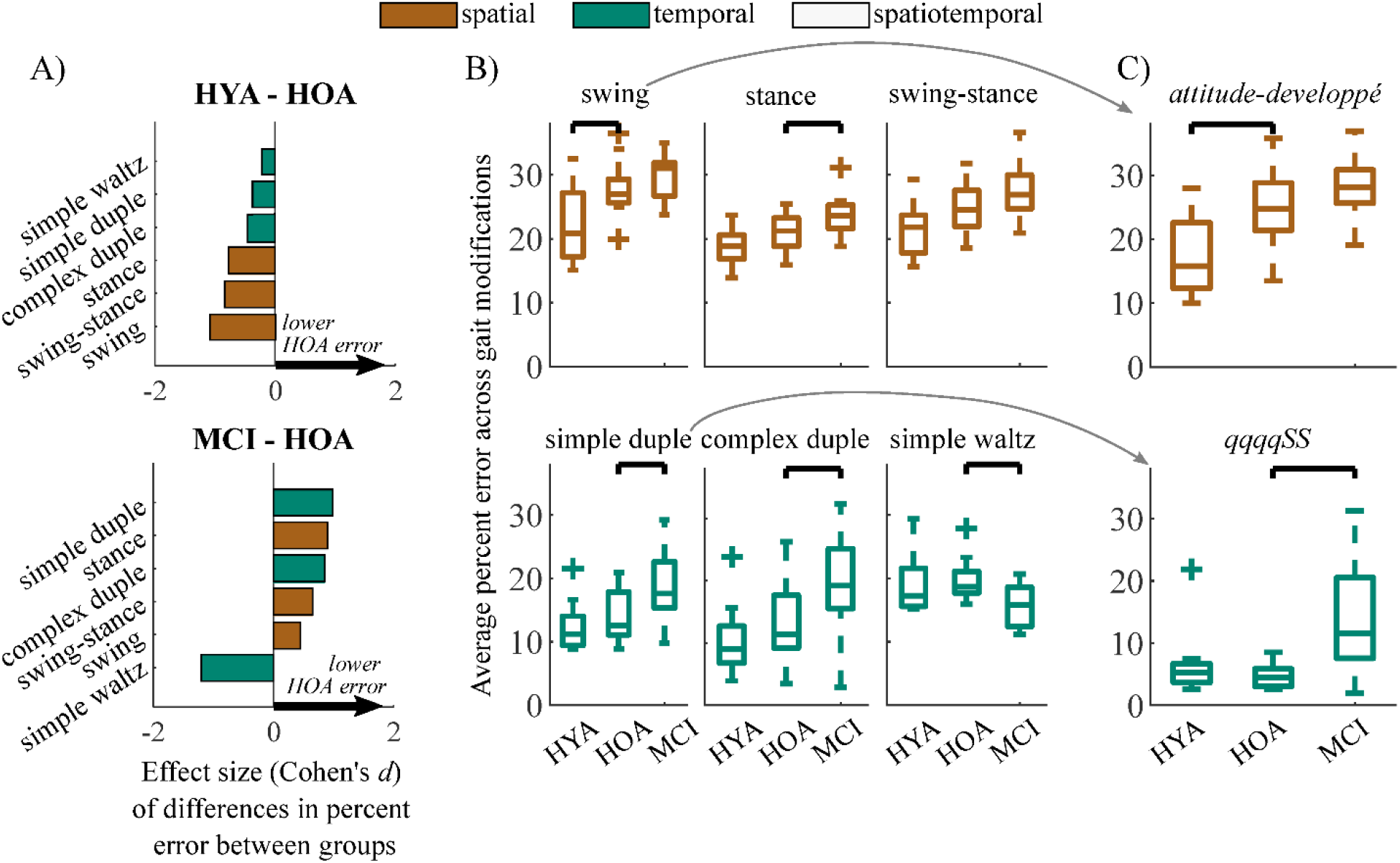
A) Effect sizes of differences in performance (Cohen’s *d*) between HYA and HOA groups (top) and HOA and MCI groups (bottom) in each gait modification subclass. Spatial and temporal modifications are denoted in orange and green, respectively. Positive values (arrows) imply that the HOA group exhibited lower error on gait modifications compared to either group. B) Boxplots showing distributions of gait modification accuracy (percent error vs. targets) for spatial (top) and temporal (bottom) gait modifications. C) Results from exemplary participants for gait modifications that elicited the largest between-group effect sizes. The *attitude-developpé* modification (top) elicited the largest difference in spatial performance between the HYA and HOA groups. The *simple duple* modification, *qqqqSS*, elicited the largest difference in temporal performance between the HOA and MCI groups but did not differentiate between the HYA and HOA groups. We did not test for differences between the HYA and MCI groups.

### RMS performance was reduced in older adults with mild cognitive impairments

Unlike differences in the HYA and HOA groups, the HOA group performed only *spatiotemporal* gait modifications significantly more accurately than the MCI group (p = 0.017; *d* = 1.3, Δ = 2.6%; **Error! Reference source not found.A**). While many participants with MCI exhibited reduced *spatial* and *temporal* modification accuracy compared to the HOA group, these differences did not reach statistical significance. *Temporal* accuracy was highly variable in both groups, with some participants with MCI performing temporal modifications more accurately than those in the HOA group. However, the HOA and MCI groups still exhibited a moderate differential effect size (p = 0.16; *d =* 0.6; Δ = 2.4%).

Differences in *spatiotemporal* modification accuracy were driven by HOA performing the *simple duple* (p = 0.012, *d* = 1.0, Δ = 4.9%), *complex duple* (p = 0.030, *d* = 0.9, Δ = 6.5%), and *stance* (p = 0.032, *d* = 0.9, Δ =2.6%) gait modifications more accurately than MCI participants (Error! Reference source not found.**A & B**). An example of group differences in *simple duple* modification accuracy is shown in Error! Reference source not found.**C**, in which participants with MCI struggled to perform the six-step temporal sequence *qqqqSS*. Surprisingly, the MCI group performed the *waltz* gait modifications significantly more accurately than the HOA group (p = 0.013, *d* = 1.2, Δ = -4.0%).

Datasheets containing clinical and demographic characteristics, effect sizes, and average differences in percent error between groups for each gait modification can be found in the *Supplemental Material*.

### Associations of motor and cognitive function with RMS performance: preliminary evidence

Exploratory correlation analyses identified non-negligible correlations in 27/29 combinations of variables tested (Table 3; asterisks designate effect sizes with cutoff values according to Cohen (1992)). The strongest correlations were between *spatial* modification performance and: Reverse Corsi Blocks product score (Pearson’s *r* = -0.50), Four-Square Step Test completion time (*r* = 0.55), and Body Position Spatial Task product score (*r* = -0.55) (**Error! Reference source not found**.). In the table, asterisks denote effect sizes for each correlation (Cohen, 1992). *Temporal* gait modifications exhibited at most moderate correlations with clinical assessments: Reverse Corsi Blocks product score (*r* = -0.31), Four-Square Step Test completion time (*r* = 0.29), and Body Position Spatial Task product score (*r* = -0.28). *Spatiotemporal* gait modifications exhibited the strongest correlations with the Four-Square Step Test completion time (*r* = 0.59), and Body Position Spatial Task product score (*r* = -0.69).

**Table 1:**
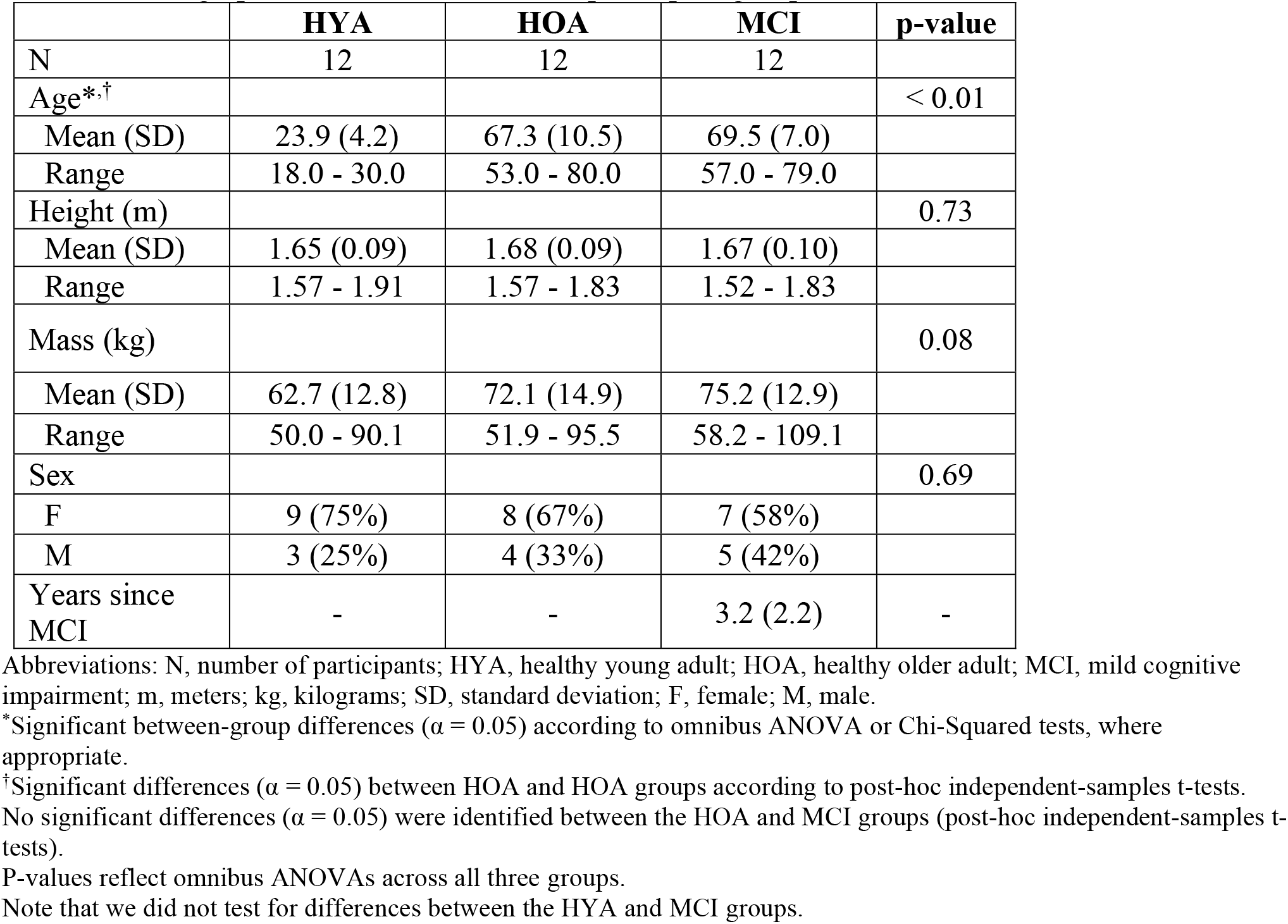
Demographic characteristics of each participant group.

|  | **HYA** | **HOA** | **MCI** | **p-value** |
| --- | --- | --- | --- | --- |
| N | 12 | 12 | 12 |  |
| Age<sup>*,†</sup> |  |  |  | < 0.01 |
| Mean (SD) | 23.9 (4.2) | 67.3 (10.5) | 69.5 (7.0) |  |
| Range | 18.0 - 30.0 | 53.0 - 80.0 | 57.0 - 79.0 |  |
| Height (m) |  |  |  | 0.73 |
| Mean (SD) | 1.65 (0.09) | 1.68 (0.09) | 1.67 (0.10) |  |
| Range | 1.57 - 1.91 | 1.57 - 1.83 | 1.52 - 1.83 |  |
| Mass (kg) |  |  |  | 0.08 |
| Mean (SD) | 62.7 (12.8) | 72.1 (14.9) | 75.2 (12.9) |  |
| Range | 50.0 - 90.1 | 51.9 - 95.5 | 58.2 - 109.1 |  |
| Sex |  |  |  | 0.69 |
| F | 9 (75%) | 8 (67%) | 7 (58%) |  |
| M | 3 (25%) | 4 (33%) | 5 (42%) |  |
| Years since MCI | - | - | 3.2 (2.2) | - |

Abbreviations: N, number of participants; HYA, healthy young adult; HOA, healthy older adult; MCI, mild cognitive impairment; m, meters; kg, kilograms; SD, standard deviation; F, female; M, male.
<sup>\*</sup>Significant between-group differences ( $\alpha = 0.05$ ) according to omnibus ANOVA or Chi-Squared tests, where appropriate.
<sup>†</sup>Significant differences ( $\alpha = 0.05$ ) between HOA and HOA groups according to post-hoc independent-samples t-tests.
No significant differences ( $\alpha = 0.05$ ) were identified between the HOA and MCI groups (post-hoc independent-samples t-tests).
P-values reflect omnibus ANOVAs across all three groups.
Note that we did not test for differences between the HYA and MCI groups.

**Table 2:**
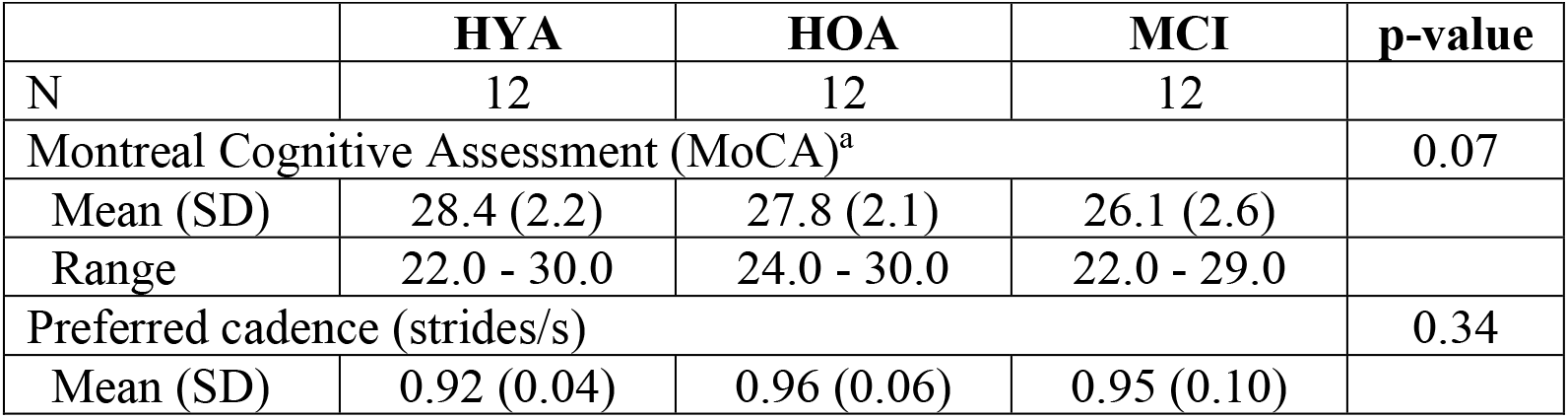
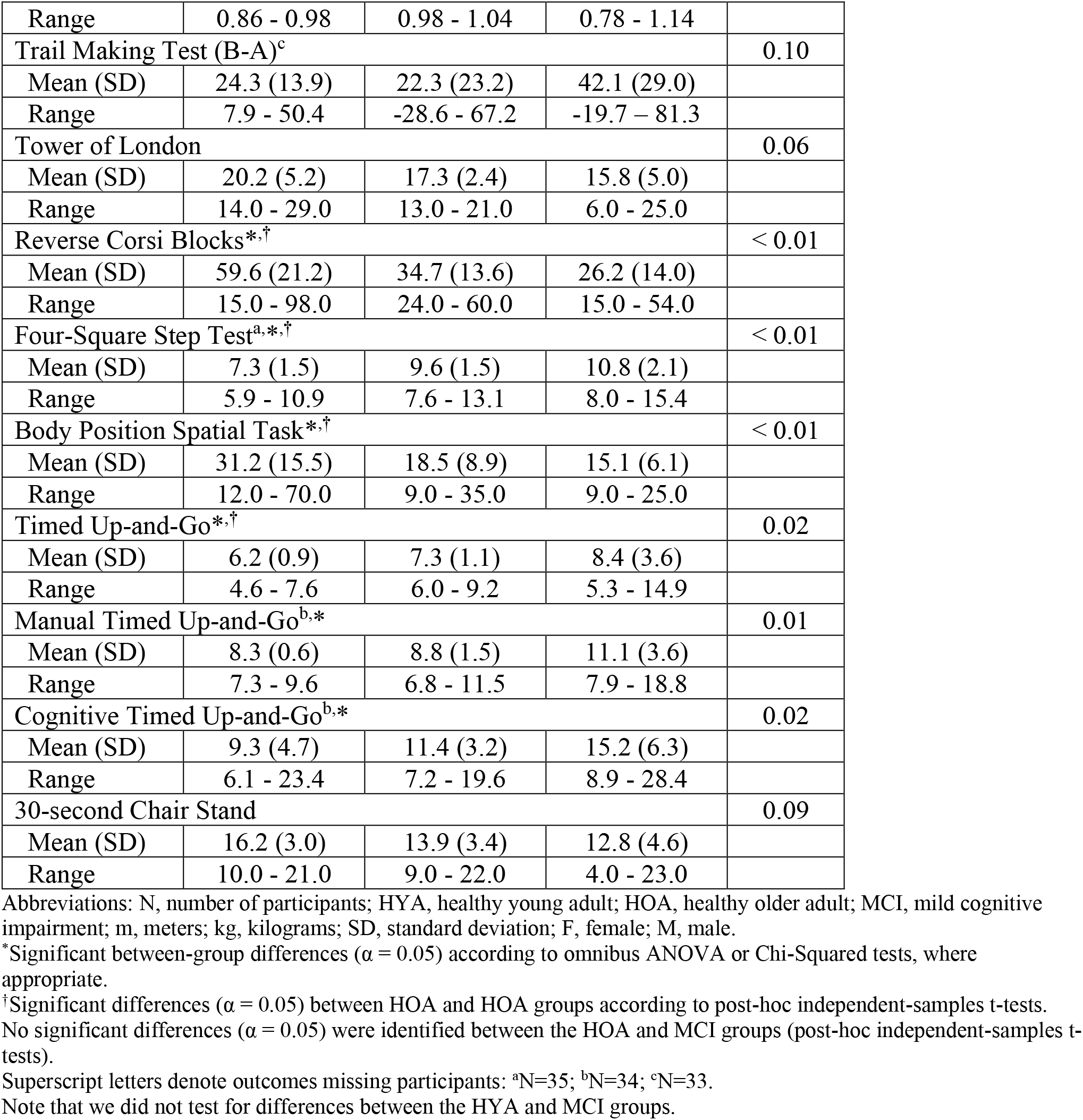
Clinical characteristics of each participant group.

**Table 3:** Correlations (Pearson’s *r*) between RMS performance in each of the three gait modification classes and clinical outcomes.

| Assessment | Gait Modification Class |  |  |
| --- | --- | --- | --- |
|  | Spatial | Temporal | Spatio-temporal |
| Trail Making Test (B-A) | 0.33** | 0.20* | 0.30** |
| Tower of London | -0.35** | -0.19* | -0.49** |
| Reverse Corsi Blocks | -0.50*** | -0.31** | -0.40** |
| Four-Square Step Test | 0.50*** | 0.29* | 0.59*** |
| Body Position Spatial Task | -0.55*** | -0.28* | -0.69*** |
| Timed Up-and-Go | 0.22* | 0.02 | 0.15* |
| Manual Timed Up-and-Go | 0.47** | 0.13* | 0.37** |
| Cognitive Timed Up-and-Go | 0.42** | -0.05 | 0.39** |
| 30-second Chair Stand | -0.25* | -0.05 | -0.27* |

\*Small effect according to Cohen (1992).
\*\*Medium effect
\*\*\*Large effect

## Discussion

This study leveraged biomechanical analysis and music theory to investigate the impacts of age and cognitive function on individuals’ abilities to flexibly modulate spatial and temporal aspects of their gait. Age-related declines in motor function (HYA vs. HOA) appear to reduce individuals’ capacities to flexibly modulate spatial aspects of gait, though cognitive deficits (MCI vs. HOA) may also contribute to reductions in this capacity. While temporal modification performance was more variable, cognitive deficits appear to reduce individuals’ abilities to modulate temporal aspects of gait. Unsurprisingly, both age-related declines in motor and cognitive function appear to reduce individuals’ abilities to flexibly modulate spatiotemporal aspects of gait, as would be done in music or dance therapy. By testing a broad set of spatial and temporal movement patterns, our novel experimental framework revealed group differences in movement performance that would not be revealed by normal walking or by a single set of gait modifications (e.g., backward walking, fast walking). Given the importance of selecting an appropriate level of challenge during rehabilitation, our findings suggest that both motor and cognitive function, as well as movement types, should be considered when designing complex movement, music-movement, or dance-based therapies for individuals with MCI as others have recommended (Guadagnoli and Lee, 2004).

### Motor and, possibly, cognitive deficits reduce the ability to accurately modulate spatial gait features

Reduced performance on spatial and spatiotemporal, but not temporal, gait modifications in HOA relative to HYA suggests that age-related declines in motor function may limit individuals’ abilities to modulate spatial components of gait. Aging is associated with reductions in strength and balance (Schloemer et al., 2017; Reimann et al., 2020). These findings are consistent with the observed large deficits in gait modification performance during *swing*-phase modifications in HOA relative to HYA, which often involved large hip ranges of motion. Additionally, a post-hoc analysis of individual gait modifications revealed that *swing* and *swing-stance* modifications comprised five of the six modifications eliciting the largest difference in performance between HOA and HYA. Reduced *swing* modification performance contrasts with performance noted during the stance modifications, which challenged strength more than balance because stance modification targets could be achieved during double-limb support. Consequently, modifications to the *swing*-phase of gait or dance steps may be critical to personalizing exercise programs based on individuals’ motor function during dance-based therapy (Bauby and Kuo, 2000; Kuo, 2007). Note that the trend toward reduced spatial modification performance in the MCI compared to the HOA group supported a potential effect of cognitive function on the ability to modulate spatial components of movement (Cohen et al., 2016; Rucco et al., 2017), which may emerge with a larger sample size.

Age-related declines in strength may impact individuals’ abilities to perform plantarflexion during *stance* modifications (Browne and Franz, 2018): multiple HOA and MCI participants were unable to generate sufficient plantarflexor moments to raise their heel during single-limb support in the *relevé and piqué* stance modifications. Plantarflexor muscle properties are altered and activation capacity is reduced in older adults, which may limit ankle torque production capacity during gait modifications (Morse et al., 2004; Conway and Franz, 2020). Dance-based movements that require plantarflexor moment generation during stance may, therefore, challenge a prominent age-related deficit in HOA and could be good targets for personalization during dance-based therapy.

Our assumption that differences in RMS performance between HYA and HOA stemmed primarily from differences in motor function was supported by worse performance on assessments of motor function (*e*.*g*., TUG, Four-Square Step Test) in HOA compared to HYA. However, some assessments involving aspects of cognitive function also differed between HYA and HOA (e.g., Body Position Spatial Task) and were correlated with spatial modification performance. Therefore, it is possible that differences in cognitive function also impact individuals’ abilities to modulate spatial aspects of gait.

### Cognitive deficits may reduce the ability to accurately modulate temporal gait features

We observed a moderate effect toward individuals with MCI exhibiting worse performance on temporal gait modifications than those in the HOA group, suggesting that cognitive deficits associated with MCI reduce the ability to flexibly modulate temporal features of gait. Such deficits correspond to the inability to entrain stepping movements to rhythms during dance classes. For example, individuals with MCI exhibited worse performance for six-step temporal sequences (vs. 2-step or 4-step sequences), particularly during the *complex duple* modifications. Accurately performing longer sequences places greater demands on working memory, which is reduced in individuals with MCI (see *Reverse Corsi Blocks Test* in **Error! Reference source not found**.) (Vandierendonck et al., 2004; Montero-Odasso et al., 2012). An alternative explanation for poor performance on some temporal modifications is that reduced motor-cognitive integration in individuals with MCI may limit individuals’ abilities to use the musical cues to adopt the proper timing for the assigned movement (Kluger et al., 1997; Hackney et al., 2017).

*Waltz* modifications represented a surprising case in which individuals with MCI performed modifications more accurately than HOA. Specifically, individuals with MCI performed the *quick-Slow* (*qS*) modification better than HOA (post-hoc analysis: Cohen’s *d* = 2.8). *Waltz* sequences were either two-or four-step, reducing their challenge to working memory compared to the six-step *Duple* sequences and possibly making these modifications easier for individuals with MCI (Montero-Odasso et al., 2012). Additionally, the three-count of *Waltz* consisted of one *quick* (*q;* 33% of the stride) and one *Slow* (*S;* 67% of the stride) step in each measure. This division of 33% and 67% of a stride is similar to the approximate percentages of the swing (40%) and stance (60%) phases during normal walking, respectively(Winter, 1984). Consequently, *Waltz* may have represented a smaller deviation from normal walking, enabling greater automaticity and reduced cortical demand than *Duple* modifications (Clark, 2015). Increased automaticity, however, would only explain a lack of deficit in *Waltz* performance in individuals with MCI. Better *Waltz* performance in MCI compared to HOA may stem from differences in music or dance experience, which represents an interesting area of future research.

Our decision to use fixed tempi for all participants is important for our interpretation of differences in temporal modification performance. Using a fixed tempo, rather than setting the tempi relative to each participant’s preferred cadence (Moumdjian et al., 2019), ensured that all participants had the same amount of time to perceive and respond to the musical cues. Personalizing the tempi to cadence would likely reduce differences in performance due to cognitive function because individuals with slower cadences would have more time to perceive the musical cues and determine motor commands to execute. Rather, we selected tempi near 100 bpm, below most participants’ preferred cadences.

Therefore, differences in temporal modification performance are likely driven by participants’ abilities to perceive cues and step in the correct patterns, rather than their physical capacity to match the tempi. This expectation is supported by HYA performing *Waltz* modifications with similar accuracy to HOA, despite stepping at a roughly 30% faster tempo. Such a fast cadence could increase demands on motor function but did not obviously reduce HYA performance. Future studies can build upon our current findings to evaluate the effect of varying tempi and movement speeds.

### Motor and cognitive deficits reduced the ability to perform spatiotemporal modifications in individuals with MCI

Large deficits in spatiotemporal performance in individuals with MCI compared to HOA were expected and further emphasize the impacts of cognitive deficits on the ability to voluntarily modulate gait biomechanics. Spatiotemporal gait modifications appear to elicit larger differences in movement performance between HOA and MCI than spatial or temporal modifications. Motor-cognition is adversely affected by MCI, and the requirement to modulate both spatial and temporal parameters affected those with MCI more than HOA (Hackney et al., 2017). Additionally, reduced set-shifting capacity in MCI compared to HOA - revealed by large differences in the A and B components of the Trail Making Task (Bowie and Harvey, 2006) - may explain reduced spatiotemporal performance in MCI. For example, individuals with MCI may struggle to identify tempos in the music while remembering gait modifications. Participants were instructed to equally prioritize the temporal or spatial components of the spatiotemporal gait modifications. However, some participants in the MCI group reported “giving up” on either the spatial or temporal components of the modifications. Future investigation into which aspects of RMS gait modifications are prioritized with age and cognitive decline may provide further insight into mechanisms impacting specific features of movement performance (Earhart, 2013).

### Potential correlates of RMS performance

Our results show that *spatial, temporal*, and *spatiotemporal* gait modifications may be correlated with assessments of visuospatial working memory (Reverse Corsi Blocks), motor-cognitive integration, balance and mobility (Four-Square Step Test), and whole body spatial memory (Body Position Spatial Task) (Dite and Temple, 2002; Vandierendonck et al., 2004; Hackney et al., 2013). These correlations suggest potential motor and cognitive contributions to spatial and temporal aspects of RMS performance and may be useful as design variables for dance-based therapy classes.

However, these findings should be considered with extreme caution because in this preliminary study, we were not able to rigorously validate the robustness of these correlations, which will be a valuable focus of future larger-sample studies. Our findings here can pilot future trials with larger sample sizes and specific hypotheses about which aspects of motor or cognitive function impact RMS performance.

### Clinical implications

Beyond advancing our understanding of how age and cognition impact the ability to flexibly modulate gait, our findings have implications for dance-based therapies for individuals with MCI. Our results showing distinct motor control deficits related to aging versus cognitive impairment inform clinical understanding of the mechanisms underlying impaired gait performance in HOA and MCI. Creative dance-based movement therapies aim to challenge both motor and cognitive function and have shown positive impacts on spatial cognition, executive function, balance, and mobility in individuals with Parkinson’s and MCI (Hackney et al., 2007a; Hackney et al., 2007b; McKee and Hackney, 2013; Zhu et al., 2020). Our results show that different temporal features of music and spatial components of movements challenge distinct motor (e.g., strength or balance) and cognitive (e.g., working memory, set-shifting, or motor-cognitive integration) features (Thom and Clare, 2011; Montero-Odasso et al., 2012; Earhart, 2013; Browne and Franz, 2018). Consequently, personalizing music and dance selections during therapy based on an individual’s motor and cognitive function may maximize therapeutic effects. Specifically, selecting music and dance parameters that optimize the level of challenge for an individual may maximize benefits while retaining motivation (Guadagnoli and Lee, 2004; Hackney et al., 2007a). For example, our results suggest that teaching spatial or temporal modifications alone may be an appropriate level of challenge for individuals with MCI, before progressing to spatiotemporal modifications (Hackney and Earhart, 2010). However, we evaluated only the immediate relationship between age-related declines in motor and cognitive function and RMS performance. With repeated practice, individuals with MCI may improve RMS performance levels to near those of HOA or HYA participants (McKee and Hackney, 2013). These learning or longitudinal effects are beyond the scope of this study but are interesting areas of future research that could inform personalized design and data-driven progression of dance-based and music-based movement therapies.

### Limitations

The following limitations constrain the generalizability of our findings. First, our modest sample size of twelve participants per group lacked the statistical power to identify all differences in movement performance, particularly during temporal gait modifications. Nonetheless, our data revealed differences in movement performance for spatial and spatiotemporal gait modifications, and showed moderate differential effect sizes (HOA vs. MCI) during temporal modification performance.

Secondly, while protocol administrators verified that participants understood each gait modification similarly before collecting data, understanding may have improved over the first few trials, potentially improving performance in later trials. However, we used a block randomization strategy to account for changes in comprehension of the movements.

Our findings should be viewed in the context of a broader set of constructs – including and beyond the motor and cognitive constraints studied here – that could impact movement performance during movement therapy. The specific gait modifications selected, sensory stimuli (music type or tempo). walking speed, or even individuals’ relationships to music and dance could impact RMS performance. For example, music with an easier-to-perceive downbeat may alter the observed relationships between cognitive function and spatiotemporal RMS performance in individuals with lesser music experience. Our group-level analyses across diverse movement classes provide an early step toward understanding how dance-based RMS challenge specific aspects of motor and cognitive function. Larger sample sizes would enable the discovery of specific factors underlying RMS performance in individuals with MCI to be discovered from a broader set of constructs affecting movement.

## Conclusions

We developed and implemented an innovative RMS gait modifications protocol to evaluate the impacts of age-related declines in motor and cognitive function on the ability to accurately modulate spatial and temporal features of gait. By improving our understanding of motor and cognitive impacts on the ability to modulate gait, this study represents an early step toward systematically personalized dance-based movement therapies. Furthermore, this study lays the foundation for future evaluation of the ability of RMS to 1) help diagnose cognitive-motor impairments by providing insight into mechanisms of movement performance deficits and 2) track intervention efficacy by quantifying changes in the accuracy of spatial and temporal features of gait modifications following therapy.

Future studies identifying clinical and mechanistic predictors of RMS performance will advance our ability to provide quantitative metrics that assist clinicians in selecting movement therapy parameters to maximize treatment efficacy.

## Supporting information

MovementLibrary

Table_Outcomes

TableStatistics

DataDictionary

Table_ClinicalScores

Table_Demographics

## Author Contributions

MCR, MEH, JLM, LE, and TMK obtained funding for the study. MEH, TMK, LE, MCR, and AS defined the study design and data collection protocol. MCR, AS, and KC were skilled assessors who performed data collection. MEH, JLM, TK, LE, and MCR supervised the study and planned data analysis. MCR and JLM performed data analysis. MCR drafted the original manuscript. All authors provided intellectual critique and manuscript revisions.

## Acknowledgments

We thank the participation of our volunteers, and T. Prusin and C. Carroll-Sauer for their assistance in data collection and participant recruitment.

## Data Availability Statement

The original contributions presented in the study are included in the supplementary material, further inquiries can be directed to the corresponding author.

