## Supplemental - S1 for "Motor and cognitive deficits limit the ability to flexibly modulate spatiotemporal gait features in older adults with mild cognitive impairment"

### 1 Description

The datasheets included as *.csv* files contain participant demographics and clinical characteristics, outcomes for each gait modification, and results of statistical analyses.

#### 1.1 Files

1. *MovementLibrary.csv* contains individual gait modifications that were combined to produce the RMS sequences used in this study and listed in *DataDictionary.csv*.
    a. Movement Class specifies a spatial or temporal modification. These combine to produce spatiotemporal modifications
    b. Movement Subclass specifies the subclass of movement: Spatial, Temporal, Simple Duple, Complex Duple, Waltz
    c. Movement specifies the movement name, which is reflected in each of the gait modifications in DataDictionary.csv.
    d. The remaining fields denote target variables and values for each movement.
        i. Spatial modification targets:
            1. Hip, knee, ankle fields have units of Degrees.
        ii. Temporal modification targets:
            1. Tempo_HYA_bpm specifies the music tempo for HYA participants in beats-per-minute. Divide by 60 (seconds/minute) and multiple by 2 (strides/step) to estimate the corresponding cadence in strides-per-second.
            2. Tempo_HOAMCI_bpm specifies the music tempo for HOA and MCI participants in beats-per-minute.
            3. Step_pattern_beatsPerStep specifies how many beats spanned each step in the sequence. In the Movement field, & = 0.5 beats, q = 1 beat, S = 2 beats.

#### 1.2 Nomenclature

*The following abbreviations and shorthand are used in the datasheets*

HYA: Healthy young adult, participants denoted as *MST0XX*

HOA: Healthy older adult, participants denoted as *MST1XX*

MCI: Mild cognitive impairment, participants denoted as *MST2XX*

RMSL Rhythmic movement sequence

SW: Spatial gait modification in only the swing-phase of gait

ST: Spatial gait modification in only the stance-phase of gait

SWST: Spatial gait modifications in both the swing- and stance-phases

SimpleDuple: Simple duple subgroup in temporal gait modifications

ComplexDuple: Complex duple subgroup in temporal gait modifications

Waltz: Waltz subgroup in temporal gait modifications

&: A step that should take one half-beat in temporal gait modifications

q: A step that should take one beat in temporal gait modifications

S: A step that should take two beats in temporal gait modifications

NaN: Trials/assessments not recorded for a participant

### 1.3 Notes

1. Each spatial gait modification used two modifications (e.g., the swing-stance modification involving the Battement swing and Piqué stance modifications would be denoted, *SWST_battementPiqué*.
